# ARDitox: platform for the prediction of TCRs potential off-target binding

**DOI:** 10.1101/2023.04.11.536336

**Authors:** Victor Murcia Pienkowski, Tamara Boschert, Piotr Skoczylas, Anna Sanecka-Duin, Maciej Jasiński, Bartłomiej Król-Józaga, Giovanni Mazzocco, Sławomir Stachura, Lukas Bunse, Jan Kaczmarczyk, Edward W Green, Agnieszka Blum

## Abstract

Cellular immunotherapies, such as those utilizing T lymphocytes expressing native or engineered T cell receptors (TCRs), have already demonstrated therapeutic efficacy. However, some high-affinity TCRs have also proved to be fatal due to off-target immunotoxicity. This process occurs when the immune system acts against epitopes found on both tumor cells and healthy tissues. Moreover, some TCRs can be cross-reactive to epitopes with highly dissimilar sequences. To address this issue, we developed ARDitox, a novel *in silico* method based on computational immunology and artificial intelligence (AI) for predicting and analyzing potential off-target binding. We tested the performance of ARDitox *in silico* on different cases found in the literature where TCRs were used to target cancer-related antigens, as well as on a set of TCRs targeting a viral epitope. ARDitox was able to identify previously reported cross-reactive epitopes in line with the data available in the literature. In addition, we investigated a TCR targeting an HLA-A*02:01-restricted immunodominant epitope from the glioblastoma-associated antigen NLGN4X, identifying a cross-reactive ADH1A epitope that would not be detected in murine models. In conclusion, our *in silico* approach is a powerful tool that identifies potential off-target epitopes, complementing preclinical studies in developing safer cell therapies targeting tumor(- associated) antigens.

## Introduction

Recent advances in the field of immunotherapy are changing the landscape of available treatments for cancer, particularly for hematological malignancies and some solid tumors ^1^. Among the most promising are adoptive cell therapies using T cells engineered to express native or engineered TCRs ^2^. While much of the field has focussed on identifying high quality tumor-specific target epitopes against which to develop TCRs, less attention has been given to the issue of predicting off-target immunotoxicity ^3^, in which T cells recognise and kill healthy cells expressing epitopes similar to the tumor(-associated) antigen target.

T cell immunosurveillance consists of TCR scanning of short peptides presented on the cell surface by receptors called human leukocyte antigens (HLA) ^4^. Importantly, only a fraction of all peptides from the human proteome is presented on the HLA. In a multi-step process, a peptide is loaded on the HLA and exported to the cell surface ^5^. One TCR can interact with multiple peptide-HLA (pHLA). TCR promiscuity is an intrinsic and necessary property of antigen recognition by T cells as the diversity of peptides presented on the cell surface is much higher than the number of TCRs in the human body ^6^. As a consequence a TCR targeting a tumor-associated epitope, can interact with an off-target epitope (OTE) presented on healthy tissue - particularly when using non-autologous TCRs that have not passed thymic selection. Furthermore, the risk of potential immunotoxicity may be increased for the TCRs engineered to enhance pHLA-TCR affinity ^3^.

Off-target toxicity has proven to represent a considerable safety liability for patients adoptively transferred with TCR-transgenic T cells as it caused death of four people in two independent clinical trials ^3,7^. Importantly, the sequences of the target epitope and the OTE do not have to be very similar. The clinically relevant cases of cross-reactivity showed that peptides with only 5 identical amino acids to the target peptide can cause off-target toxicity ^8^. Unfortunately, using experimental methods to test all possible off-targets is costly and time-consuming thus has to be limited to a restricted subspace of potential OTEs.

Leveraging recent advances in computational immunology and AI can augment these efforts, increasing the number and safety of available treatments. To this end, we introduce **ARDitox** - a novel method for predicting OTEs for candidate therapeutic TCRs.

## Methods

### The ARDitox pipeline

The pipeline of our method consists of 5 consecutive steps (Figure 1) as discussed in the following sections.

**Figure 1.**
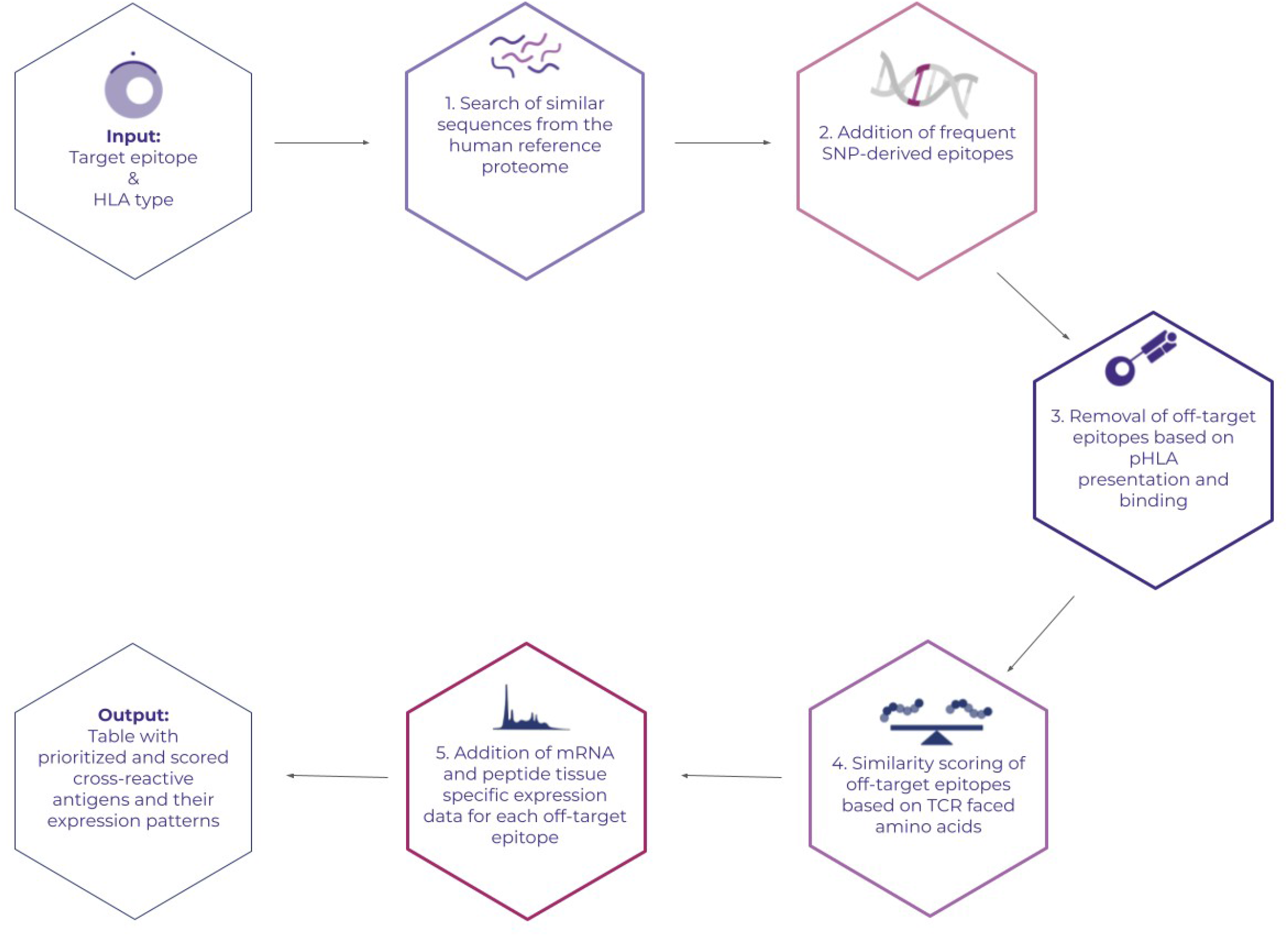
Workflow of ARDitox

#### 1. Identification of all putative off-target sequences

ARDitox takes as an input a target epitope from 8 to 11 amino acids long and its corresponding HLA type. Firstly, the algorithm identifies all epitopes that have at least 5 amino acids shared with the target epitope. Importantly, only OTEs of the same length are taken into account as it is considered rare for epitopes of different lengths to bind to the same TCR ^9^. This step generates a large number of putative OTEs, e.g. for an 8 amino acid target epitope there are combinatorially 459360 possible putative OTEs according to 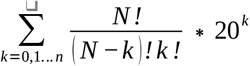 where k is the number of substituted amino acids, n is the maximum number of substituted amino acids so 5 of them are shared with the target epitope and N is the length of the epitope. Each of these putative OTEs is then checked for presence in the human reference proteome (UniProt-https://www.uniprot.org/proteomes/UP000005640) ^10^.

#### 2. Addition of Single nucleotide variant epitopes

Single nucleotide polymorphisms (SNPs) are a major source of OTEs as a given human genome contains ∼7000 nonsynonymous germline SNPs ^11^. As such, SNPs can be a major source of novel off-target sites in TCR-based therapies. Unfortunately, SNPs are not included in the UniProt proteome reference. ARDitox introduces frequent nonsynonymous mutations (frequency >1% in at least one studied population) from gnomAD database ^12^ (https://gnomad.broadinstitute.org/). Each identified SNP occurring in the sequence corresponding to the OTEs is taken into account through the addition of an additional putative OTE. This step further increases the number of OTEs to be analyzed, making the algorithm more sensitive.

#### 3. Selection of Presented epitopes

The previous steps generate an enormous number of potential OTEs, of which only a fraction will be presented on an HLA class I allele. In order to limit the number of putative OTEs to the presented ones, we use an in-house developed presentation model called ARDisplay ^13^. The presentation model is based on artificial neural networks trained using results of mass-spectroscopy experiments e.g Sarkizova et al. ^14^. Overall, it generates predictions for canonical class I HLA (i.e., A, B, and C). Only OTEs that have a probability of being presented >50% (ARDisplay) and binding affinity <2000 nM (MHCflurry^15^) proceed to the next steps.

#### 4. Off-target epitopes ranking

In the target epitope, amino acids in different positions can interact with the HLA and with the TCR. In order for these interactions to occur, the physico-chemical properties of the amino acids at certain positions must remain similar ^16^. In this step, we establish the positions of TCR-faced residues, depending on the HLA type the epitope binds to. The positions are based on literature e.g. Bijen et al. ^17^ and database search (https://www.iedb.org/) containing information, which residues are crucial for TCR binding. The comparison between the target epitope and putative OTE is performed on the TCR-facing residues using selected amino acids physico-chemical properties obtained from https://www.genome.jp/. All the matrices containing these properties were linearized into vectors. The distance between each target epitope *e* and the putative OTE *p* can be computed according to the following norm:

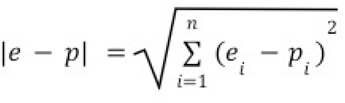

where *e*_*i*_ and *p*_i_ represent the i-th physico-chemical property of the amino acids from peptides e and p, and n is equal to the target peptide length.

#### 5. mRNA and peptide tissue specific expression

mRNA expression in TPMs (GTEx) ^18^ and protein expression level (Human Protein Atlas) ^19^ of the putative OTEs are added in order to identify tissues sensitive to off-target toxicity.

## In vitro validation

### mRNA electroporation for TCR expression

TCRs were ordered from TWIST Biosciences in custom vectors. In vitro transcribed RNA was generated using T7 Scribe Standard RNA IVT Kit (Biozym #150404). RNA transfection was performed by electroporation using the 4D-Nucleofector electroporation system (Lonza, program CL-120, solution SE).

### Cell culture

Jurkat T cells (Leibnitz Institute DSMZ #ACC282) were cultured in RPMI 1640 Medium (Gibco, #61870143) with 10% heat-inactivated FBS (PAN-Biotech) and 1% penicillin-streptomycin (Capricorn Scientific #PS-B). Commercially available EBV-immortalized HLA-A*02:01 expressing BOLETH cells were used as antigen presenting cells. The BOLETH cells were cultured with 50mM β-mercaptoethanol, 1mM sodium pyruvate (Thermo Fisher Scientific #11360039), and 1X MEM Non-Essential Amino Acids (Gibco, #11140035). BOLETH and T cells were plated in 1:3 ratio in a round-bottom 96-well plate and the respective peptides were added at a final concentration of 10µM. As a positive control, Jurkat T cells were plated in a well pre-coated with CD28/CD3 monoclonal antibodies at a 1:400 dilution. After 16h of co-culture T cell activation was assessed.

### Flow Cytometry

Cells were diluted and washed with FACS buffer (PBS + 2mM EDTA + 2% FBS) and centrifuged at 300g for 5min. Supernatant was removed and cells resuspended in 50µl staining solution containing diluted antibodies as follows:

| Specificity | Fluorophore | Clone | Supplier | Product number | Dilution |
| --- | --- | --- | --- | --- | --- |
| CD2 | PerCP/Cy5 | RPA-2.10 | BioLegend | 300216 | 1:200 |
| CD3 | Unlabeled | OKT3 | BioLegend | 317325 | 1:400 |
| CD20 | PacificBlue | 2H7 | BioLegend | 302320 | 1:200 |
| CD28 | Unlabeled | E18 | BioLegend | 122022 | 1:400 |
| CD69 | PE/Cy7 | FN50 | BioLegend | 310912 | 1:200 |
| Mouse-TCR-β | PE | H57-597 | BioLegend | 109208 | 1:200 |

Following 20min incubation at 4°C cells were washed twice with FACS buffer, pelleted (300g for 5min), and resuspended in 100µL FACS buffer.

### Dataset preparation

We selected three groups of peptides presented on either HLA-A*02:01 or HLA-A*01:01 for the evaluation of our methodology: i) Tumor Associated Antigen (TAA) epitopes, ii) known immunodominant viral epitopes, and iii) epitopes derived from frameshift mutations. TAA and virus epitopes were acquired from IEDB ^20^. Frameshift derived epitopes were obtained from a library of Neo Open Reading Frame peptides (Supplementary table 1) ^21^. We used ARDisplay to predict frameshift epitopes presented by HLA-A*02:01, and downsampled the dataset to 16 epitopes. Importantly, we randomly selected 16 TAAs presented on HLA-A*02:01 in order to compare TAA vs frameshift epitopes on equally abundant groups.

## Results

ARDitox estimates the safety of the putative OTEs through a Safety score that can vary from 0 to 14. A score close to 0 means that the putative OTE and the target epitope are predicted to have an almost identical pHLA:TCR interaction, and it is highly probable for cross-reactive binding to occur.

## Known cross-reactive epitopes

We tested ARDitox on 5 cases of TCRs targeting TAA epitopes ^3,7,22-24^ and on 1 case of a set of T cells targeting a viral peptide ^25^ described below case by case. The target epitopes and their respective cross-reactive epitopes are presented in Table 1.

**Table1.**
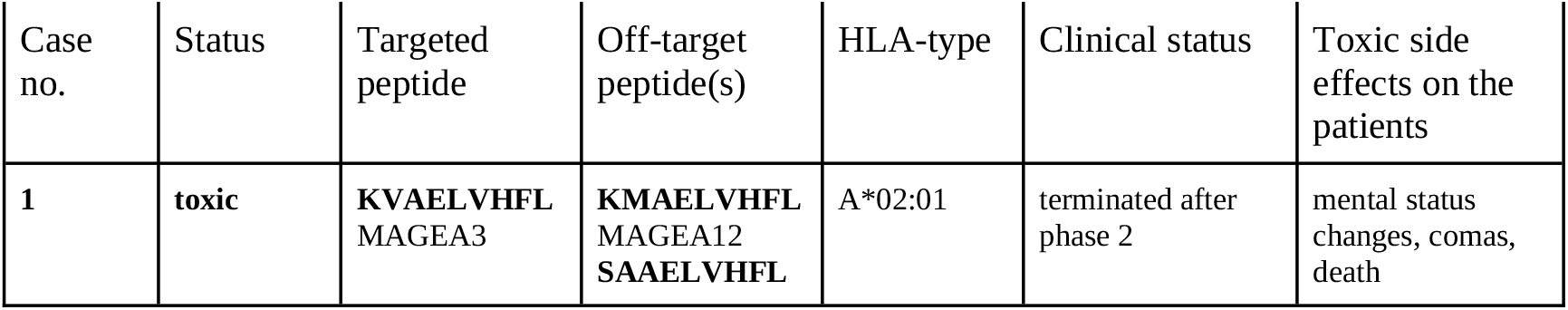
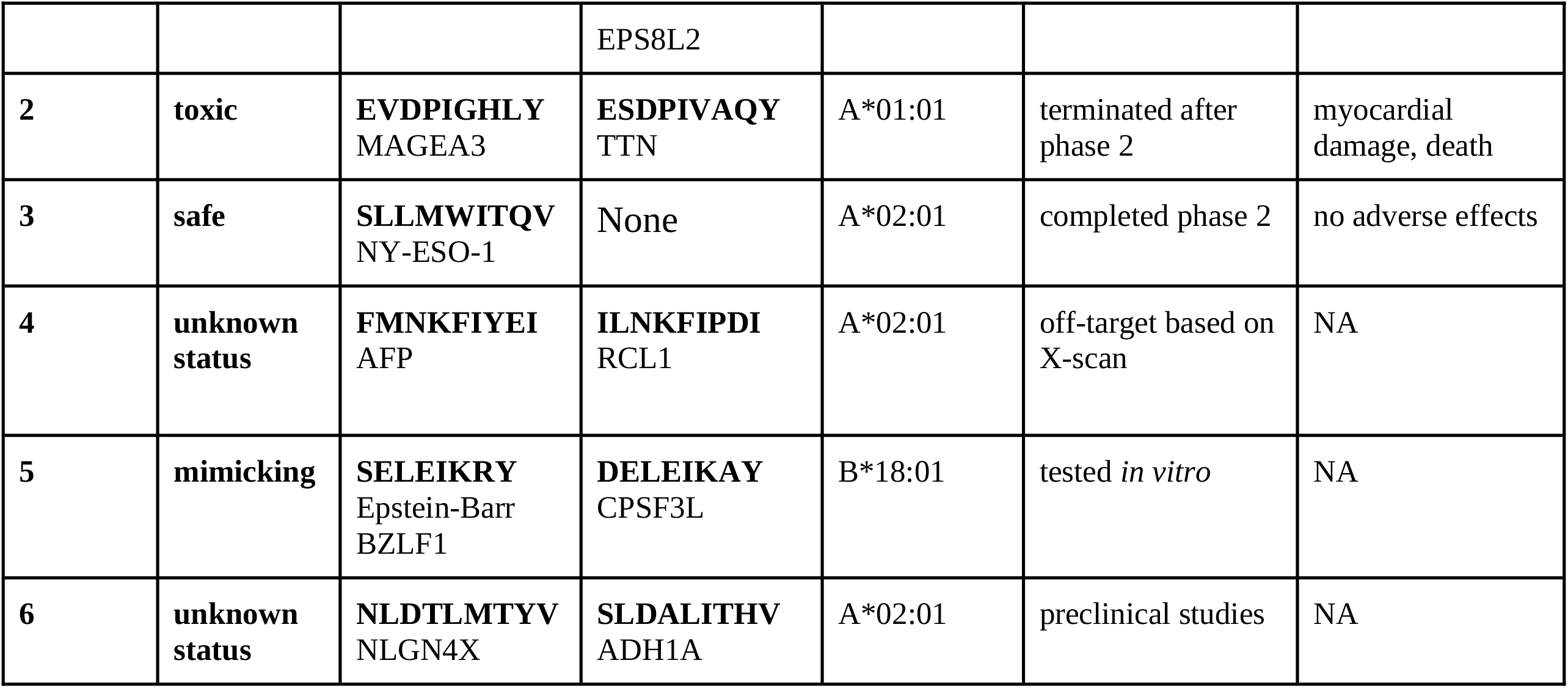
Target epitopes used for ARDitox validation, together with their properties and status obtained in previous studies.

### Case 1

Morgan et al. ^7^ described a cell therapy treating nine HLA-A*02:01 positive patients with metastatic cancer expressing MAGEA3 (KVAELVHFL) and MAGEA12. Three patients developed severe neurotoxic side effects, two of which proved fatal. The authors showed that MAGEA12 and to a lower degree MAGEA1, MAGEA8, and MAGEA9 expressed in the brain are probably responsible for the toxicity effects. A re-evaluation of the OTEs that could be responsible for the neurotoxicity of this TCR pointed to EPS8L2 (SAAELVHFL) as a possible culprit of the observed site effects ^26^.

ARDitox, when applied to the target epitope KVAELVHFL, identified 294 putative OTEs (Supplementary table 2) that both shared at least 5 amino acids with the target and had a high probability of being presented on the cell surface. Among them, 24 OTEs had a safety score <3.0 (Figure 2A), including putative OTEs derived from *MAGEA12, MAGEA8, MAGEA9*, and *EPS8L2*.

**Figure 2.**
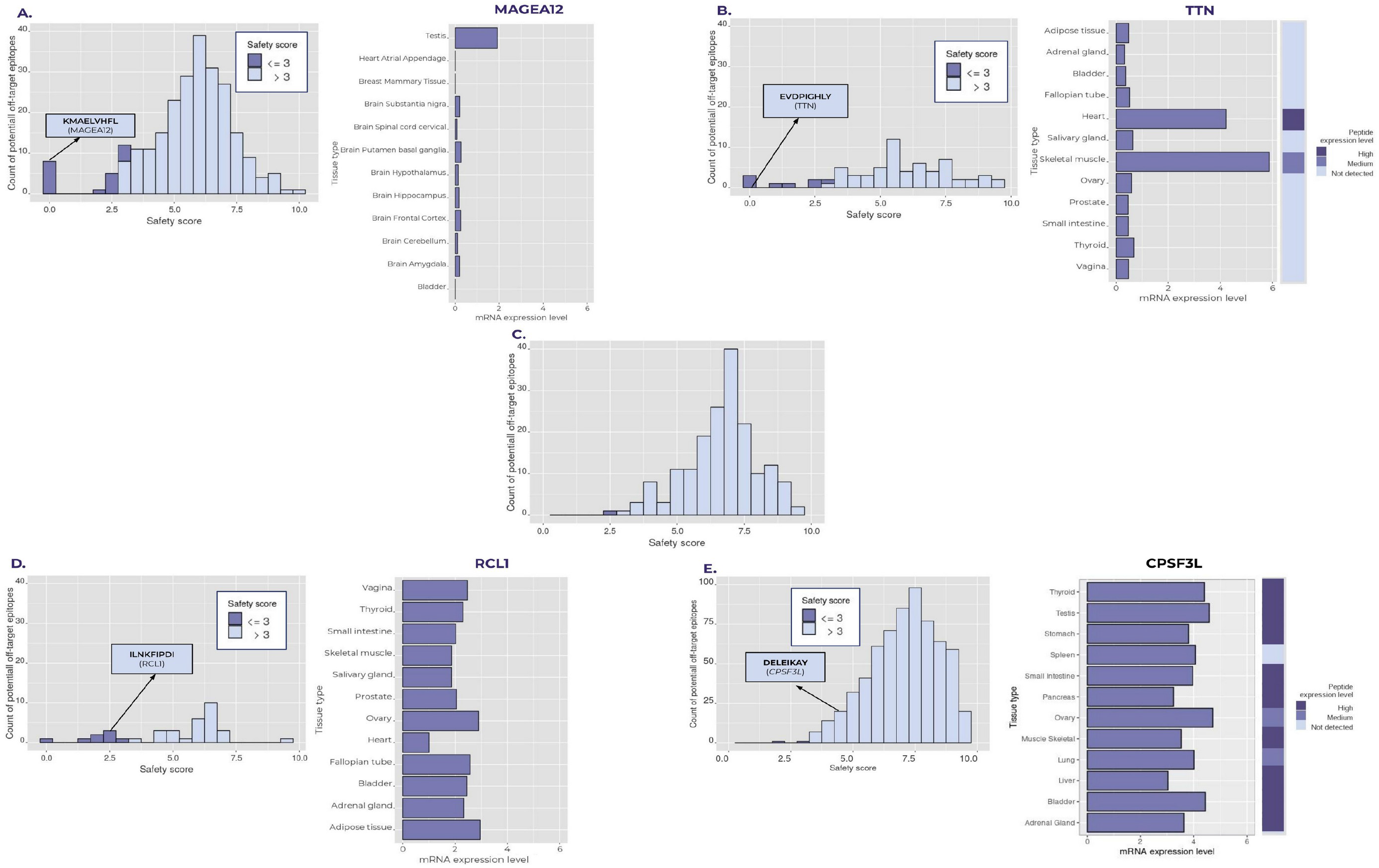
Distribution of ARDitox safety scores for putative off-target epitopes (A) case 1-MAGEA3 (KVAELVHFL), (B) case 2-MAGEA3 (EVDPIGHLY), (C) case 3-NY-ESO-1 (SLLMWITQV), (D) case 4-AFP (FMNKFIYEI) (F) case 5-Epstein-Barr *BZLF1* (SELEIKRY). If available mRNA (x=axis in logTPM) and peptide expression values of the genes that gave rise to the off-target epitope are presented.

### Case 2

Linette et al. ^3^ described a case of off-target toxicity in a clinical study conducted on four patients with myeloma and melanoma. The immunotherapy was directed against a MAGEA3 epitope (EVDPIGHLY) presented on HLA-A*01:01. Two patients that received the therapy developed cardiogenic shock and as a result died within the next few days. Only after performing the experiments on cultured beating cardiomyocytes, a titin (TTN) epitope was identified to be responsible for the toxicity effects.

ARDitox identifies 84 potential OTEs, among which nine have a safety score <3.0 (Figure 2B). Importantly, the epitope originating from TTN was one of the top hits with a safety score of 0 (Supplementary table 2).

### Case 3

Stadtmaueret al. ^22^ performed a clinical study on 25 high-risk multiple myeloma patients using T cells engineered against NY-ESO-1 (SLLMWITQV), a cancer-testis antigen with expression in multiple types of cancer. The TCR-based cancer immunotherapies targeting NY-ESO-1 are considered as the most promising ones.

This SLLMWITQV epitope is presented on HLA-A*02:01. For that pHLA combination, ARDitox finds 203 putative OTEs. Out of all OTEs only a single epitope derived from LRBA (FLLMFIKQL) has a safety score <3.0 (Figure 2F, Supplementary table 2).

### Case 4

Cai et al. ^23^ tested preclinically engineered T cells against AFP epitope (**FMNKFIYEI**) presented on HLA-A*02:01. Based on an *in vitro* X-scan experiment, two OTEs that could activate the T cell were identified: ENPP1_436_ and RCL1_215_. However, ENPP1_436_ is neither processed nor presented on the human cells.

Results from ARDitox point out 39 putative OTEs for this target. Eight of them are characterized by a safety score <3.0. Among them, an experimentally identified epitope (X-scan); RCL1_215_ is graded by ARDitox with a safety score of 2.47 indicating its high off-target binding potential (Figure 2C, Supplementary table 2).

### Case 5

Rist et al. ^25^ showed that a high proportion of CD8+ T cells against an EBV epitope from BZLF1 (SELEIKRY) protein presented on HLA-B*18:01 cross-react with a human off-target epitope CPSF3L (DELEIKAY). The authors hypothesized that BZLF1 is an example of molecular mimicry.

ARDitox identified 661 putative OTEs for SELEIKRY. Among them, DELEIKAY was found with a safety score of 3.94 and was one of the top 15 cross-reactive peptides (Figure 2D, Supplementary table 2).

### Case 6

Lastly, we validated ARDitox predictions in *in vitro* experiments. We chose NLGN4X_131–139_ (NLDTLMTYV) that has been reported as promising recurrent TAA in glioblastoma ^24^. We performed prospective safety analysis to evaluate the suitability of this epitope as a cell therapy target. This epitope is presented on HLA-A*02:01. For that pHLA combination, ARDitox found a single putative OTE: NLGN4Y with a safety score <3.0, and 16 additional putative OTEs with a safety score <5 (Figure 3A, Supplementary table 2). All the epitopes with a safety score <5 were further verified *in vitro* as described in the Methods section. This resulted in the identification of an OTE with a score of 4.87 (SLDALITHV; *ADH1A*) that weakly activated the examined TCR (Figure 3B-D).

**Figure 3.**
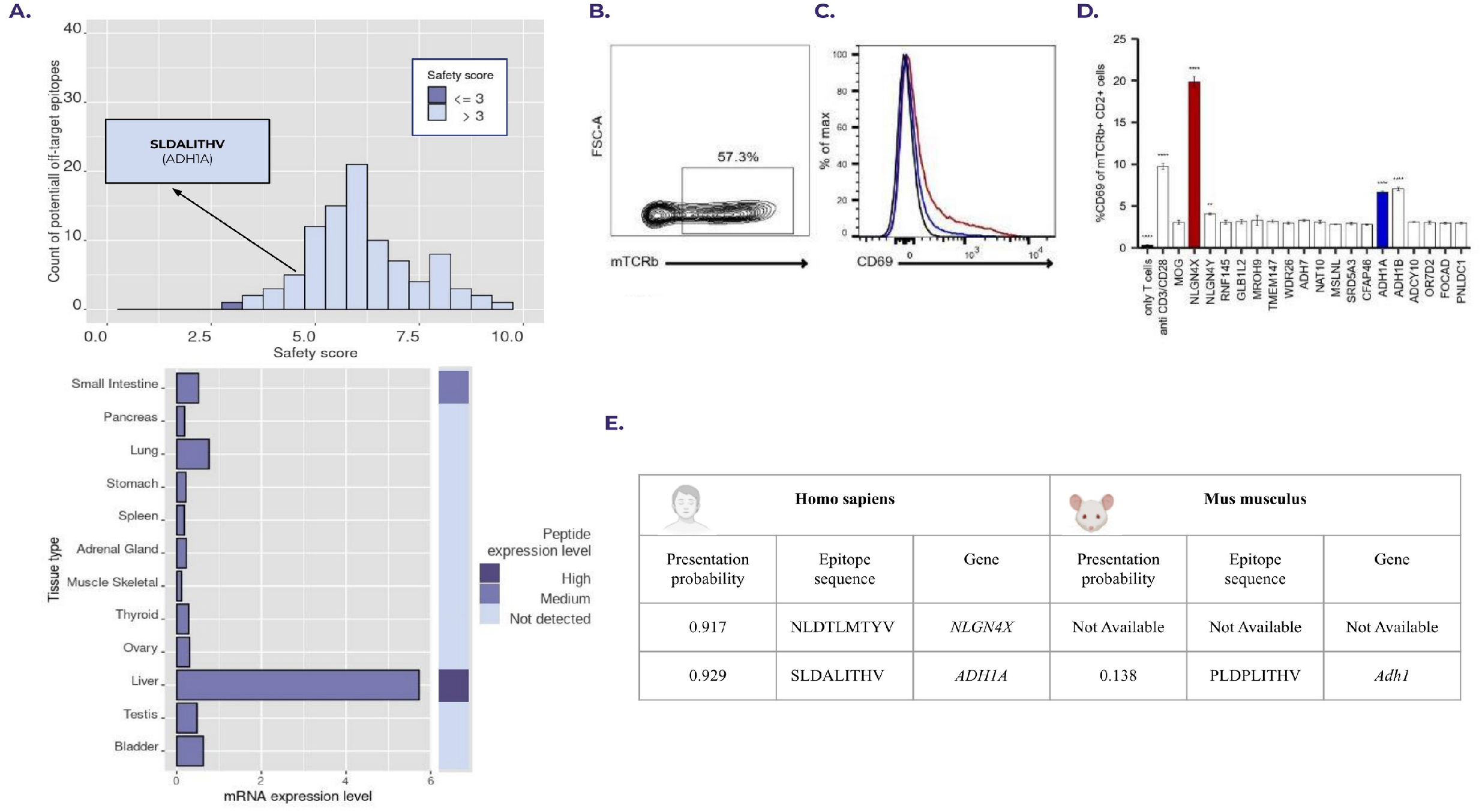
Results for case 6-NLGN4X (NLDTLMTYV) (A) Distribution of ARDitox safety scores for putative OTEs and expression plots for ADH1A OTE. (B) Representative flow cytometry density plot depicting transfection efficiency of murine TCR (mTCRb) in Jurkat cells. (C) Histogram depicting CD69 of NLGN4X TCR Jurkat T cells co-cultured with NLGN4X (red) or ADH1A peptide (blue)-pulsed presenter cells are overlayed on only T cell control (black). (D) CD69 levels of mTCRb+ Jurkat T cells (CD2+) after co-culture with presenter cells loaded with the indicated putative OTEs. Represented is the mean with SD of n=3 technical replicates. Significance calculated with one-way ANOVA in comparison to the Myelin Oligodendrocyte Glycoprotein (MOG) control peptide. (E) Table showing the lack of presentation of epitope PLDPLITHV derived from *Adh1* a mouse ortholog gene of *ADH1A*. In this particular case the mouse model would omit the ADH1A OTE.

### OTE trends

#### TAA epitopes vs Virus epitopes

We employed ARDitox to analyze 148 epitopes from TAA and viruses presented on HLA-A*02:01 or HLA-A*01:01. The number of putative OTEs for TAA epitopes was higher (31854) than OTEs of viral origin (22279). The t-test conducted on the safety scores suggests a significant difference between the distributions (mean TAA=7.39, mean Virus=7.62, p-value<2.2e-16). On the other hand, Cohen’s-d (−0.1514) ^27^ suggests a rather negligible difference. The above values, together with the similar shapes of distributions of TAA and Virus OTEs (Figure 4) suggest that the distribution of safety scores is comparable between the two groups. However, if only OTEs with safety scores below 3 are considered, we see a 6.4-fold enrichment of TAA vs viral epitopes.

**Figure 4.**
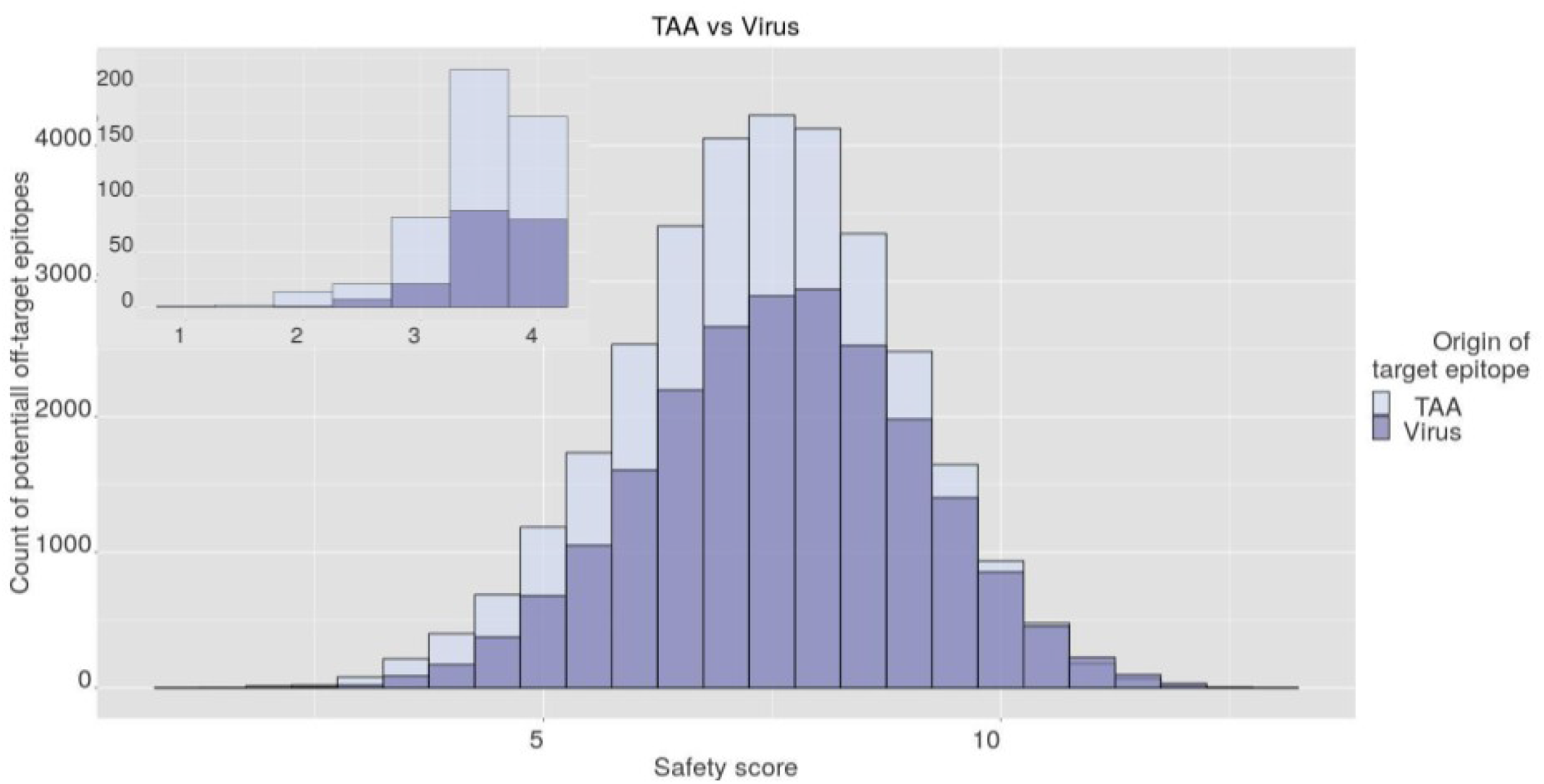
Distribution of ARDitox safety scores of OTEs from 148 TAA epitopes and 148 virus epitopes presented on HLA-A*02:01 or HLA-A*01:01

#### TAA epitopes vs frameshift epitopes

Lastly, ARDitox was tested on 16 frameshift epitopes and 16 randomly subsampled TAA peptides, predicted by our model to be presented on HLA-A*02:01. Importantly, only 336 putative OTEs were identified for the frameshift-derived fragments, while 3911 putative OTEs were found in the TAA group. The t-test conducted on the safety scores was significant with a p-value=2.544e-06 (mean TAA=7.21, mean Frameshift=7.67), while Cohen’s d value was low (0.28). Interestingly, no OTE with a score below 3 was found for targets derived from frameshift variants (Figure 5).

**Figure 5.**
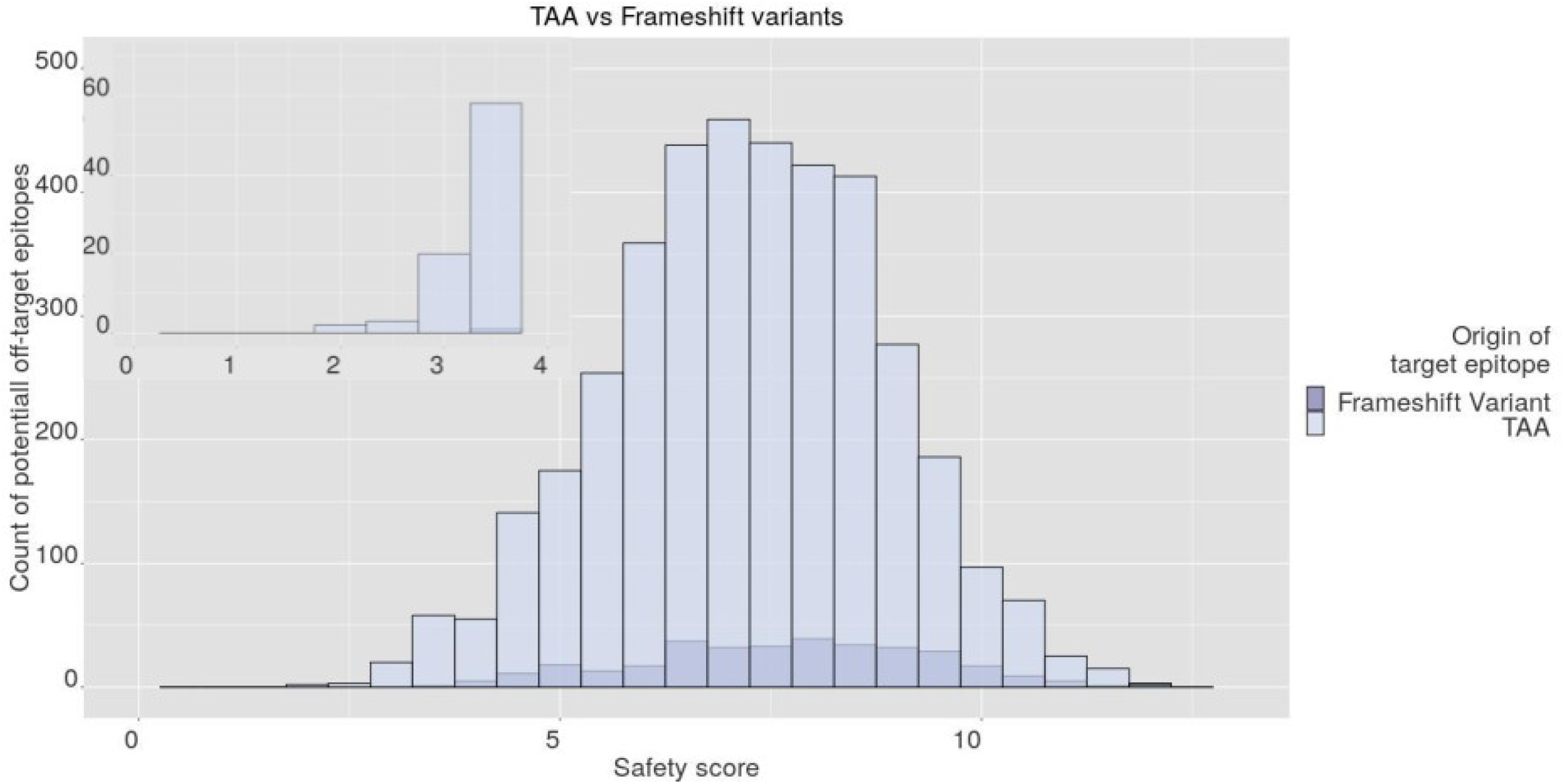
Distribution of ARDitox safety scores of OTEs from 16 TAA epitopes and 16 frameshift epitopes presented on HLA-A*02:01

## Discussion

Cellular cancer immunotherapy is on the way to constitute a major pillar for cancer treatment, next to surgery, radiotherapy, and chemotherapy ^28^. One of the main issues that need to be addressed is the risk of adverse effects caused by off-target toxicity ^29^. To help mitigate this problem we have introduced **ARDitox** - a novel method for analyzing potential cross-reactivity for a given pHLA, that includes the identification of OTEs whose sequence differs significantly from the targeted epitope.

So far very few computational algorithms for predicting off-target binding or off-target toxicity in TCR-based cellular cancer immunotherapies have been proposed. Similarly to ARDitox, Expitope ^30^ and iVax ^16^ approaches start with querying human proteome for peptides homologous to the target with a predefined number of mismatches allowed. However, iVax does not consider the presentation of recognized putative OTEs in the process of ranking them and Expitope estimates peptide presentation on HLA with a proxy by the combination of the proteasomal cleavage probability and transporter associated with antigen processing as well as HLA binding. This means that ARDitox is the first available method leveraging a model for predicting peptide presentation by HLA molecules and unlike Expitope includes recognition of the TCR facing residues in order to evaluate the safety of the cross-reactive epitopes. Overall, ARDitox is a novel approach to OTEs identification with unique pipeline that includes an in-house AI trained presentation model, a unique scoring function focused on physico-chemical properties of the TCR-facing residues and an extended search of peptides derived from frequent mutations.

We tested ARDitox on four clinically validated cases of TCRs targeting TAA epitopes, one TCR in a preclinical stage and one virus epitope as well as on two datasets: i) TAA vs Virus epitopes and ii) TAA vs frameshift derived epitopes. In all analyzed cases with reported side effects, ARDitox correctly identified the OTEs that caused the toxicity in the treated patients (Table 2). What is more, in case 3, where no risky OTEs were found experimentally, ARDitox found only one cross-reactive epitope with a safety score <3. In case 5 we successfully identified an OTE that might lead to autoimmunity as a result of molecular mimicry after EBV infection, showing ARDitox usefulness in e.g. viral vaccine design that takes into account potential of molecular mimicry. Lastly, In case 6 we experimentally found the ADH1A epitope, an off-target that would not be identified in mouse models due to lack of presentation of the orthologue epitope derived from *Adh1* (PLDPLITHV) making it impossible to find this OTE with mouse models (Figure 3E). Importantly, ADH1A has a safety score close to 5, which corresponds with the weak binding of the TCR (Figure 3 C-D). Early identification of this OTE is valuable as additional safety measures can be taken to ensure that activation of the T cell against ADH1A will not occur during clinical trials.

**Table 2.**
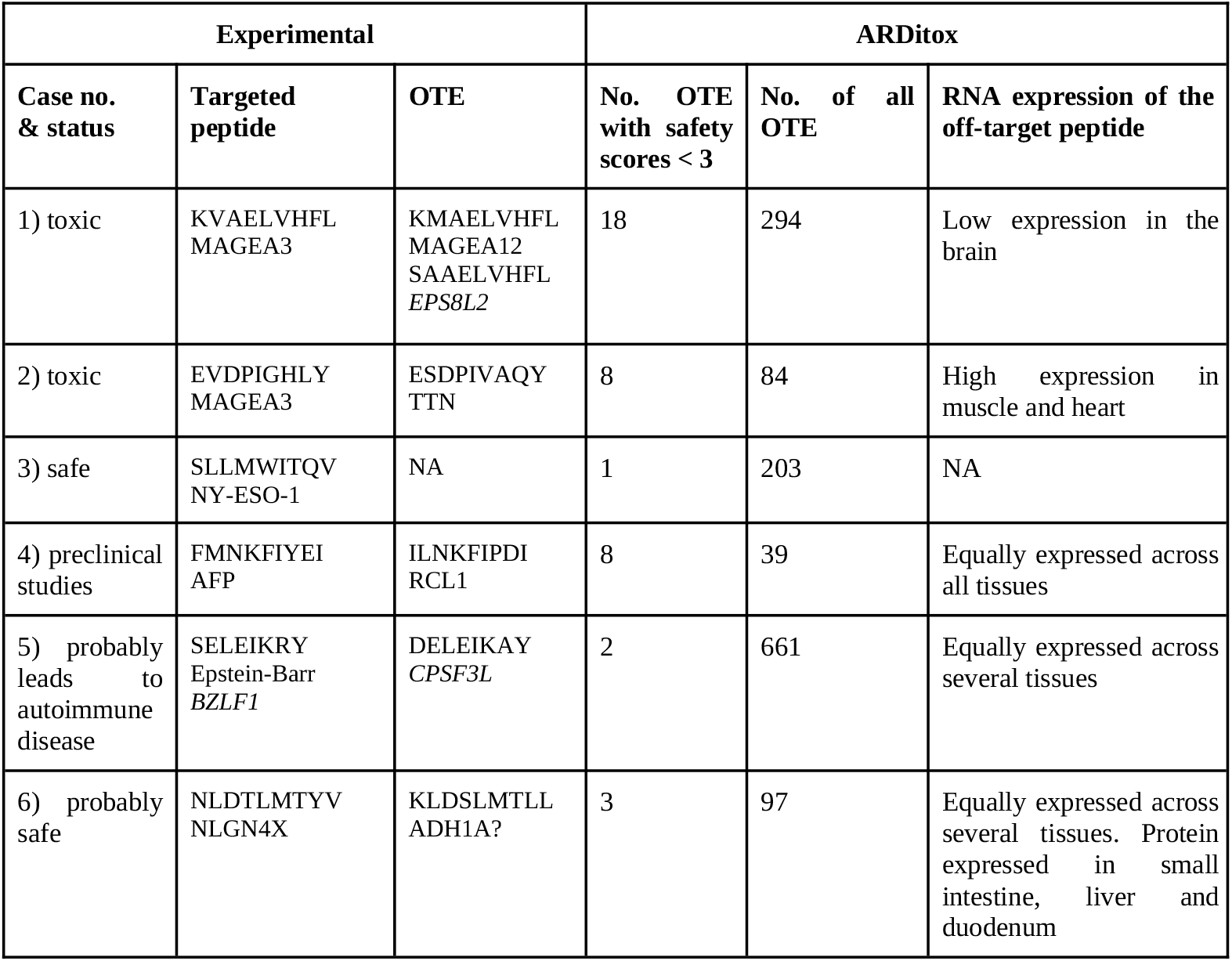
Estimation of targeted peptide toxicity by ARDitox based on three main variables

When using ARDitox to assess the risk of the therapy, we recommend checking three variables presented by the software: i) Number of all OTEs, ii) Number of OTEs with a safety score <3, and iii) expression of OTEs with a safety score <3 (Table 2). Furthermore, we highly recommend testing *in vitro* all OTEs with a score <5 as some weak TCR off-target interactions (e.g. case 5 and 6) might have negative consequences for the patient’s health. As an example to illustrate the weights of the variables mentioned above we will use *case 2* with an OTE from TTN protein.. The number of all putative OTEs in *case 2* should be considered moderate as only 84 were identified (Table 2). Based on this variable, the TCR against this target epitope seems to be very promising. However, when the distribution of the safety scores is verified, it turns out that ∼10% of the putative OTEs have a score <3. The cross-reactivity of each of these OTEs should be checked experimentally because low safety scores indicate that the TCR may bind to both the target and the OTE in a similar fashion. Lastly, checking the expression status of each putative OTE with a safety score <3 should indicate which tissue types are of particular interest for the experimental verification. This would have been important in *case 2*, as during the preclinical *in vitro* studies no toxicity towards the tested heart muscle cell line was found, as TTN protein is expressed only in contracting cardiac myocytes ^4^. Identifying TTN as OTE upfront could enforce adding appropriate cell lines to the test panel. The potential shortcoming of our model is that currently, for some less frequent HLA-types, incorrect amino acids may be scored as TCR faced. However, this problem is minor as it occurs only for HLA-types that have generally low frequency. Furthermore, it can be mitigated as more data regarding TCR-faced amino acid positions for rare HLAs becomes available.

Lastly, in order to assess the effectiveness of the proteome search for OTEs and the proposed scoring methodology, we compared the analysis performed on TAAs and viral epitopes. As expected, due to the evolutionary distance between the tested peptides the differences between the amount of OTEs derived from viruses with a score <3 are much lower than in TAAs epitopes, indicating that additional care should be taken when assessing the risk of the off-targets with a safety score <3. Furthermore, we observed the same trend for frameshift mutations. We saw no frameshift OTEs with a score <3 and observed a very high ratio of frameshift OTEs vs TAA OTEs with a score <5. This confirmed our expectations as frameshifts give rise to multiple, out-of-frame, random protein products that differ significantly from non-mutated self-proteome indicating that frameshift neoepitopes can be safe and promising alternatives to TAA epitopes in TCR therapies ^31^.

In conclusion, we have developed a method for the identification of potential off-target binding in cellular immunotherapies. Our tool, named **ARDitox**, takes into account: (i) peptide processing, (ii) pHLA binding, (iii) pHLA presentation probability, (iv) determination and similarity of TCR-faced amino acids, (v) frequent variants as a source of OTEs, and (vi) gene mRNA and protein expression levels. Most importantly, by testing the method on several cases we proved its efficiency in the identification of OTEs.

## Disclosure of potential conflicts of interest

No potential conflicts of interest were disclosed. ARDitox is under *European Patent Office* (*EPO*) revision number EP22461636.7

## Availability of data and material

All datasets are available in the supplementary files. The source code for ARDitox is not available.

## Authors’ contributions

AB and EWG contributed equally to the study. VMP, ASD, TB, JK, GM and EWG wrote the manuscript. TB, EWG, LB performed the experiments. VMP, PS, BKJ, GM, MJ and SS developed ARDitox and performed the analysis of the tool. VMP, AB and JK planned the functionalities of ARDitox. All authors read and approved the manuscript.

## Funding

The research was supported by grant entitled “AImmune - Artificial Intelligence Algorithm for Identification of Immunogenic Neoepitopes of Cancer to Predict and Boost Patient’s Response to Immunotherapies”, co-financed by the Regional Operational Programme for the Małopolska Region 2014-2020, grant no: RPMP.01.02.01-12-0301/17.

## Competing interests

None declared.

## Notes

### Competing Interest Statement

The authors have declared no competing interest.

